# Proper Conditional Analysis in the Presence of Missing Data Identified Novel Independently Associated Low Frequency Variants in Nicotine Dependence Genes

**DOI:** 10.1101/222695

**Authors:** Bibo Jiang, Sai Chen, Yu Jiang, Mengzhen Liu, William G. Iacono, John K. Hewitt, John E. Hokanson, Kenneth Krauter, Markku Laakso, Kevin W. Li, Sharon M. Lutz, Matthew McGue, Daniel McGuire, Anita Pandit, Gregory Zajac, Michael Boehnke, Goncalo R. Abecasis, Scott I. Vrieze, Xiaowei Zhan, Dajiang J. Liu

**Affiliations:** Department of Public Health Sciences, Penn State College of Medicine, Hershey, PA, 17033.; Center of Statistical Genetics, Department of Biostatistics, University of Michigan, Ann Arbor, MI, 48109.; Institute for Behavioral Genetics, University of Colorado Boulder.; Department of Psychology, University of Minnesota, Minneapolis, MN 55454.; Department of Epidemiology, School of Public Health, University of Colorado Denver, Aurora, Colorado 80045.; Department of Medicine, University of Eastern Finland and Kuopio University Hospital, Kuopio, Finland.; Department of Biostatistics and Informatics, University of Colorado, Anschutz Medical Campus, Aurora, CO.; Department of Clinical Science, Quantitative Biomedical Research Center, University of Texas Southwestern Medical Center, Dallas, TX 75390.

## Abstract

Meta-analysis of genetic association studies increases sample size and the power for mapping complex traits. Existing methods are mostly developed for datasets without missing values. In practice, genotype imputation is not always effective, e.g. when targeted genotyping/sequencing assays are used or when the un-typed genetic variant is rare. Therefore, contributed summary statistics often contain missing values. Naïve extensions of existing methods either replace missing summary statistics with 0 or discard studies with missing data. These approaches can bias genetic effect estimates and lead to seriously inflated type-I or II errors in conditional analysis, which is a critical tool for identifying independently associated variants.

To address this challenge and complement imputation methods, we developed a method to combine summary statistics across participating studies and consistently estimate joint effects, even when the contributed summary statistics contain large amount of missing values. Based on this estimator, we propose a score statistic we call PCBS (partial correlation based score statistic) for conditional analysis of single-variant and gene-level associations. Through extensive analysis of simulated and real data, we showed that the new method produces well-calibrated type-I errors and is substantially more powerful than existing approaches. We applied the proposed approach to analyze the *CHRNA5-CHRNB4-CHRNA3* locus in a large-scale meta-analysis for cigarettes-per-day. Using the new method, we identified three novel variants, independent of known association signals, which were otherwise missed by alternative methods. Together, the phenotypic variance explained by these variants is .46%, improving that of previously reported associations by 17%. These findings illustrate the extent of locus allelic heterogeneity and can help pinpoint causal variants.

**AUTHOR SUMMARY:** It is of great interest to estimate the joint and conditional effects of multiple correlated variants from large scale meta-analysis, in order to fine map causal variants and understand the genetic architecture for complex traits. The contributed summary statistics from participating studies in a meta-analysis often contain missing values, as the imputation methods are not often effective, especially when the underlying genetic variant is rare or the participating studies use targeted genotyping array that is not suitable for imputation. Existing meta-analysis methods do not properly handle missing data, and can incorrectly estimate correlations between score statistics. As a result, they can produce highly biased estimates of joint effects and highly inflated type-I errors for conditional analysis, which will in turn result in overestimated phenotypic variance explained and incorrect identification of causal variants. We systematically evaluated this bias and proposed a novel partial correlation based score statistic. The new statistic has valid type-I errors for conditional analysis and much higher power than the existing methods, even when the contributed summary statistics in the meta-analysis contain a large fraction of missing values. We expect this method to be highly useful in the sequencing age for complex trait genetics.

## INTRODUCTION

Meta-analysis has become a critical tool for genetic association studies in human genetics. Meta-analysis increases sample sizes, empowers association studies, and has led to many exciting discoveries in the past decade [1-5]. Many of these genetic discoveries have informed new biology, provided novel clinical insights [6, 7], and led to novel therapeutic drug targets [8, 9]. Conditional meta-analysis has been a key component for these studies, which is useful to distinguish novel association signals from shadows of known association signals and to pinpoint causal variants.

Existing methods for conditional meta-analysis were proposed based upon the assumptions that summary association statistics from all variant sites are measured and shared in meta-analysis. Yet, in practice, summary association statistics from contributing studies often contain missing values, possibly due to the use of different genotyping arrays, sequencing capture assays, or quality control filters applied by each participating cohort. While genotype imputation is an effective approach to fill in missing genotype data for participating cohorts, many scenarios may preclude accurate genotype imputation. For example, a targeted genotyping array/sequencing assay (e.g. exome array) may not provide sufficient genome-wide coverage for imputation. In addition, it is challenging to impute low frequency variants even with the highest quality reference panels, and imputed genotypes of low quality are often filtered out. It is therefore important to properly perform meta-analysis in the presence of missing values from contributed summary statistics.

When contributed summary statistics from participating studies contain missing values, a simple strategy for marginal (or unconditional) analysis is to replace missing summary statistics with zero (REPLACE0), which is their expected value under the null hypothesis [2, 3]. This method yields valid type I errors for marginal association analysis, and is more powerful than strategies that discard studies with missing data (DISCARD). Taking this simple approach for conditional analysis, however, is problematic. The genetic variants at conditioned sites likely have non-zero effects. Replacing missing summary data with zero will bias the genetic effect estimates at conditioned variant sites, and can lead to highly inflated type I errors for conditional analysis (see RESULTS). On the other hand, discarding studies with missing summary statistics at conditioned variant sites will give valid type I errors, but at the cost of reduced power. No satisfactory solution has been described for conditional analysis when summary statistics from contributing studies have missing values.

To overcome the limitations of existing methods, we developed an improved conditional meta-analysis method that borrows strength across multiple participating studies and consistently estimates the partial variance-covariance matrices between genotypes and phenotypes. The new method is a partial correlation based score statistic (PCBS), which yields correct type I errors in the presence of missing data and is much more powerful than aforementioned simple modifications of existing methods. Interestingly, when missingness only occurs at the variant sites that we condition on, the new method PCBS has comparable power to the analysis of the complete dataset with no missing data.

We applied PCBS (together with existing methods) to a large meta-analysis on cigarettes per day (CPD). Applying the new method, we identified three new independently associated variants at the known CPD locus, *CHRNA5-CHRNB4-CHRNA3*, independent from previously reported GWAS signals. Together, these variants explained .46% of the trait variance, which improved the phenotype variance explained by previously reported GWAS hits (0.34%) by 17%. The “chip” heritability for CPD was estimated to be 5.4% [10], so the newly identified associations explained around 10% of the “chip” heritability.

To maximize the impact of the proposed method, we implemented it in our widely used software tools RAREMETAL[11] and R package rareMETALS and made them publically available. (<u>https://genome.sph.umich.edu/wiki/Rare_Variant_Analysis_and_Meta-Analysis</u>). We expect these methods to play an important role in sequence-based genetic studies and lead to important genetic discoveries in large datasets.

## MATERIALS AND METHODS

In this section, we first review the standard meta-analysis methods for single variant and gene-level association tests when analyzing datasets without missing summary statistics from contributing studies. We then illustrate the limitations of the methods for conditional analysis and describe the new method PCBS for valid and powerful conditional analysis in the presence of missing summary statistics from contributing studies.

### Overview of Meta-analysis Methods

We denote the genotype for individual *i* at variant site *j* in study *k* as *G_ijk_* which can take values of 0,1 or 2, representing the number of the minor (or alternative) alleles in the locus. When the genotypes are imputed or generated from low pass sequencing studies, genotype dosage can be used in association analysis. In this case, *G_ijk_* will be the expected number of minor (or alternative) allele counts. We denote the non-genotype covariates as *Z_ik_*, which includes a vector of 1’s to incorporate the intercept in the model. Single variant association can be analyzed in a regression model: 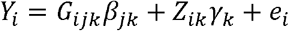. The score statistic for single variant association takes the form:

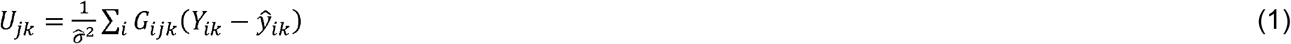

where 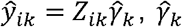, is the covariate effect, and is *σ̂* the standard deviation for the phenotype residuals estimated under the null. We denote the vector of score statistics in the region as **U_k_** = (*U*_1*k*_, …, *U_Jk_*). The variance-covariance matrix between scores statistics is equal to

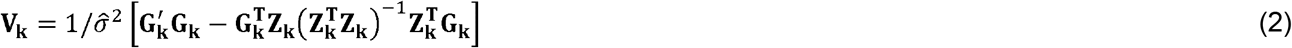

For the illustration of the method, we focused on the analysis of continuous outcomes, yet the meta-analysis and conditional meta-analysis methods work for both continuous outcomes and binary outcomes.

The meta-analysis score statistics and their covariance matrices are calculated using the Mantel-Haenszel method, i.e. 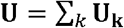 and 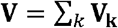. The meta-analysis statistics can be used to estimate the joint effects for variants 1,…, *J*, i.e. 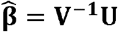.

We denote the score statistics at candidate and conditioned variant sites as **U** = (**U_G_**, **U_G^*^_**) with variance covariance matrix 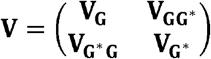

The conditional score statistic can be calculated by

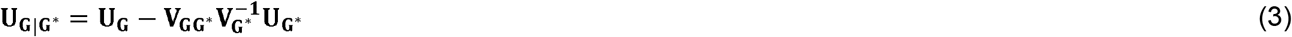

It is easy to verify the variance of the conditional score statistics is equal to

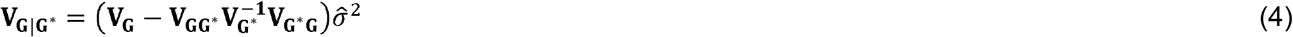

The single variant and gene-level tests in conditional analysis can be calculated based upon the conditional score statistics **U_G|G^*^_** and the covariance matrix **V_G|G^*^_**. Details are provided in **Text S1**.

### Naïve Methods In the Presence of Missing Summary Statistics

When the contributed summary association statistics from participating studies contain missing values, the REPLACE0 method replaces missing summary statistics with zero. We denote the resulting statistics as **U^0^** and **V^0^**. To mathematically describe this method, we define an indicator variable *M_jk_*, which takes value 1 if the summary statistics at site *j* in study *k* is measured and 0 if missing. The meta-analysis score statistic is calculated by

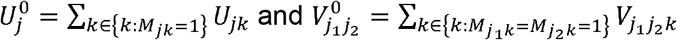

We proved in **S1 Text** that replacing missing summary association statistics with zero will bias the genetic effect estimate, i.e. 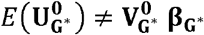. As a consequence, under the null hypothesis that the candidate variant is not associated with the phenotype, the expectation of the conditional score statistics is not equal to 0, i.e. 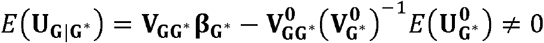. The type I error for conditional analysis can be highly inflated.

An alternative approach we call DISCARD, is to remove studies with missing summary statistics and only use studies with complete data. The meta-analysis score statistics under this analysis strategy are given by:

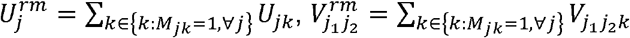

An obvious limitation of the DISCARD method is that it may result in the removal of a large number of studies and a significant loss of power.

### Partial Correlation Based Score Statistics (PCBS)

Reviewing formulae (3) and (4), note that the conditional score statistics and their variances only depend on the partial variance-covariance matrix between the phenotypes and the genotypes after the adjustment of covariates. The key idea underlying our approach is to derive a consistent estimator for the partial covariances in the presence of missing summary statistics and to use it for unbiased conditional analysis.

In statistics, to calculate the partial covariance between random variables *G_jk_* and *Y_k_* adjusting for variable *Z_k_*, we first regress out covariate *Z_k_* from both *G_jk_* and *Y_k_*, and then calculate the covariance between the residuals. Specifically,

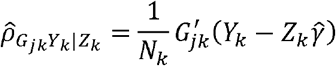

For a given study, it is easy to check that the partial covariances are scaled score statistics, i.e.

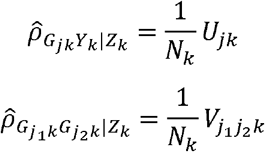

Therefore, in meta-analysis, we propose to estimate the partial covariance between genotype *G_ij_*, phenotype *Y_t_* after adjusting the covariate effect *Z_i_* using all available summary statistics:

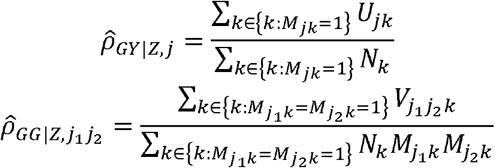

For notational convenience, we define the matrices of partial covariance as 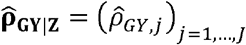 and 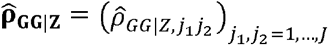. Under the fixed effect meta-analysis, we have 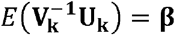 for all *k*. We showed in **S1 Text** that 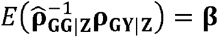. Therefore, the partial covariance matrix can be consistently estimated even in the presence of missing summary statistics.

We define partial correlation based score statistics as

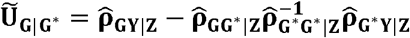

The covariances 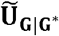 for are equal to

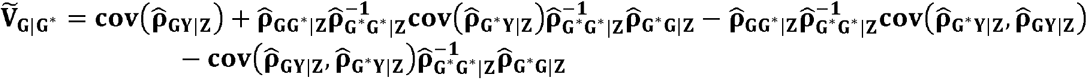

It is easy to verify that the conditional analysis using the estimator 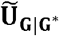 is equivalent to the standard score statistics when no missing data are present. In the presence of missing data, the partial correlation based statistic 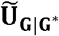 remains consistent. The conditional association analysis can be performed by replacing the standard score statistic with a partial correlation based score statistic. Details for calculating single variant and gene-level conditional association statistics can be found in **S1 Text**.

### Simulation Study

We conducted extensive simulations to evaluate the performance of PCBS as well as the two alternative approaches REPLACE0 and DISCARD. We simulated genetic data following a coalescent model that we previously used for evaluating rare variant association analysis methods[2]. The model captures an ancient population bottleneck and recent explosive population growth. Model parameters were tuned such that the site frequency spectrum and the fraction of the singletons of the simulated data match that of the real sequence data from the exome sequencing projects.

Phenotype data from each cohort were simulated according the linear model:

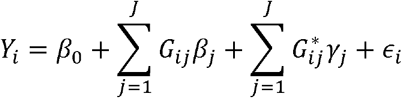

which assumes that the rare variants have additive effects on the phenotype. The genetic effects for candidate variants follow a mixture normal distribution, which accommodates the possibility that a genetic variant can be causal (with probability *c*) or non-causal (with probability 1 − *c*): 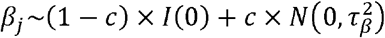. The genetic effects for the conditioned variants follow: 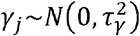.

To evaluate the influence of missing data, we randomly chose a certain fraction of the sites from each study and masked them as missing. We then applied the new method PCBS, along with DISCARD and REPLACE0 to the data to compare performance. We evaluated the type I errors and power for each approach under a variety of scenarios with different genetic effect sizes, fractions of causal variants in the gene region, and the fraction of missing data.

### Analysis of Real Data

To evaluate the effectiveness of methods in real datasets, we applied our methods to a meta-analysis of seven cohorts with a cigarettes-per-day (CPD) phenotype. Participating studies were the Minnesota Center for Twin and Family Research (MCTFR)[12-14], SardiNIA[15], <u>MET</u>abolic <u>S</u>yndrome <u>l</u>n <u>M</u>en (METSIM)[16], Genes for Good[17], COPDGene with samples of European ancestry[18], Center for Antisocial Drug Dependence (CADD)[19], and UK Biobank. Summary association statistics from the seven cohorts were generated using RVTESTS[20], and meta-analysis performed using RAREMETAL with the PCBS statistics and other competing approaches. Detailed descriptions of the cohorts are available in **S1 Text** section 4, including the methods for association analyses and the adjusted covariates.

To ensure the validity of our association analysis results, we conducted extensive quality control for the imputed genotype data. We filtered out variant sites with the imputation quality metric *R*^2^ < .7 and sites that showed large differences in allele frequencies from the imputation reference panel. Imputation dosages were used in the association analysis.

We applied iterative single variant conditional analysis to identify independent associated variants in each locus. We started by conditioning on the most significant variant from marginal association analysis. After each round of the association analysis, if the top variant remained statistically significant, we added the top variant to the set of conditioned variants, and performed an additional round of association testing. We applied three methods to analyze the data, including the partial correlation based score statistic, the method that replaces missing summary statistics with 0 and the method that discards studies with missing data. In order to examine if the low frequency variants in aggregate can be explained by the identified independently associated variants, we also performed gene-level association analysis for rare variants with MAF<5%, conditional on the identified independently associated variants.

## RESULTS

### Evaluation of Type I Error and Power of PCBS Statistics

We evaluated the type I errors for the three conditional analysis methods PCBS, REPLACE0, and DISCARD. Scenarios were considered for different combinations of the fractions of missing data, the genetic effects of the variants in the candidate gene, and the genetic effects of the conditioned variants.

First, we noted that the naïve approach REPLACE0, which replaces missing summary statistics with zero, may induce seriously inflated type I errors, under realistic patterns of linkage disequilibrium based upon our coalescent simulation. For example, when the genetic effect of the variants that we conditioned on is .05, and 50% of summary association statistics from each study were masked as missing, the type I error is 0.0025, which is 5 times the size of the significance threshold *α* = 0.0005 (**Table 1**). The type I error can be even more inflated when rate of missingness is high or when the effect sizes of the conditioned variants are large. For example, when the effect of the variant that we conditioned on is .1, and 50% of the summary association statistics from each study are masked as missing, the type I error for the naïve approach is 0.023, which is >40 times the significance thresholds. Similar inflations in the type I errors were observed for gene-level tests. The inflation in the type I errors increases with the effects of the conditioned variants and the fraction of missing data. When the conditioned variant has effect .1 and the rate of missingness is 50%, the type I errors for simple burden, SKAT and VT are .038/.040/0.032 which are up to 80-fold inflated (**Table 2**).

**Table 1:** Power and Type I Errors of Meta-analysis of Single Variant Tests in the Presence of Missing Data. Datasets were simulated according to the genetic and phenotype model described in METHODS. Metaanalysis was performed to combine 10 cohorts with 2000 individuals each. For each replicate, summary association statistics were generated, and a certain fraction of the generated summary statistics were masked as missing. Scenarios with different combinations of known variant effect, candidate variant effects and fractions of missingness were considered. Three analysis strategies were considered: 1) PCBS - partial correlation based statistics; 2) DISCARD - only analyze studies with complete summary statistics 3) REPLACE0 - replace missing summary statistics with zero. Type I errors and power were evaluated using 1 million replicates under the significance threshold of *α* = 0.0005.

| Conditioned Variant Effect | Candidate Variant Effect ($\tau_\beta$) | Fraction of Missing Data | Type I Error/Power | | | |
| --- | --- | --- | --- | --- | --- | --- |
|  |  |  | PCBS | DISCARD | REPLACE0* | Analyze the Full Dataset [Gold Standard] |
|  |  |  | Type I Error |  |  |  |
| 0.05 | 0 | 0.1 | $3.2 \times 10^{-4}$ | $3.7 \times 10^{-4}$ | $6.1 \times 10^{-4}$ | $3.7 \times 10^{-4}$ |
| 0.05 | 0 | 0.3 | $4.0 \times 10^{-4}$ | $4.5 \times 10^{-4}$ | $1.3 \times 10^{-3}$ | $3.7 \times 10^{-4}$ |
| 0.05 | 0 | 0.5 | $4.5 \times 10^{-4}$ | $1.8 \times 10^{-4}$ | $2.5 \times 10^{-3}$ | $3.7 \times 10^{-4}$ |
| 0.1 | 0 | 0.1 | $3.2 \times 10^{-4}$ | $3.7 \times 10^{-4}$ | $1.2 \times 10^{-3}$ | $3.7 \times 10^{-4}$ |
| 0.1 | 0 | 0.3 | $4.5 \times 10^{-4}$ | $4.5 \times 10^{-4}$ | $9.0 \times 10^{-3}$ | $3.7 \times 10^{-4}$ |
| 0.1 | 0 | 0.5 | $6.0 \times 10^{-4}$ | $2.6 \times 10^{-4}$ | 0.023 | $3.7 \times 10^{-4}$ |
| Power |  |  |  |  |  |  |
| 0.05 | 0.1 | 0.1 | 0.042 | 0.035 | - | 0.064 |
| 0.05 | 0.1 | 0.3 | 0.022 | 0.019 | - | 0.064 |
| 0.05 | 0.1 | 0.5 | 0.012 | $6.9 \times 10^{-3}$ | - | 0.064 |
| 0.1 | 0.1 | 0.1 | 0.046 | 0.042 | - | 0.064 |
| 0.1 | 0.1 | 0.3 | 0.022 | 0.019 | - | 0.064 |
| 0.1 | 0.1 | 0.5 | 0.013 | $6.9 \times 10^{-3}$ | - | 0.064 |
| 0.05 | 0.2 | 0.1 | 0.53 | 0.51 | - | 0.67 |
| 0.05 | 0.2 | 0.3 | 0.35 | 0.24 | - | 0.67 |
| 0.05 | 0.2 | 0.5 | 0.19 | 0.071 | - | 0.67 |
| 0.1 | 0.2 | 0.1 | 0.53 | 0.50 | - | 0.67 |
| 0.1 | 0.2 | 0.3 | 0.35 | 0.24 | - | 0.67 |
| 0.1 | 0.2 | 0.5 | 0.19 | 0.071 | - | 0.67 |

\*: For the method that replaces missing summary statistics with 0, the type I error can be severely inflated. So power was not evaluated.

**Table 2:** Power and Type I Errors of Meta-analysis of Gene-level Tests in the Presence of Missing Data and Genetic Effect Heterogeneity. Datasets were simulated according to the genetic and phenotype model described in METHODS. Within the gene region, 20% of the variant sites are deemed causal. Meta-analysis was performed to combine 10 cohorts with 2000 individuals each. For each replicate, summary association statistics were generated, and a certain fraction of the generated summary statistics were masked as missing. Scenarios with different combinations of known variant effect, candidate variant effects and fractions of missingness were considered. 1) PCBS - partial correlation based statistics; 2) DISCARD - only analyze studies with complete summary statistics 3) REPLACE0 - replace missing summary statistics with zero. To evaluate the power loss due to missing data, we also analyze the full dataset as a gold standard. Type I errors and power were evaluated for three rare variant tests (simple burden, SKAT and VT) using 1 million replicates under the significance threshold of *α* = 0.0005.

| Conditioned Variant Effect | Candidate Variant Effect ($\tau_\beta$) | Fraction of Missing Data | Type I Error/Power for Burden/SKAT/VT ($\alpha=0.0005$) | | | |
| --- | --- | --- | --- | --- | --- | --- |
|  |  |  | PCBS | DISCARD | REPLACE0* | Analyze the Full Dataset [Gold Standard] |
| Type I Error |  |  |  |  |  |  |
| 0.05 | 0 | 0.1 | $4.5 \times 10^{-4}/3.1 \times 10^{-4}/3.8 \times 10^{-4}$ | $5.2 \times 10^{-4}/5.2 \times 10^{-4}/5.9 \times 10^{-4}$ | $4.1 \times 10^{-4}/4.1 \times 10^{-4}/5.9 \times 10^{-4}$ | $4.8 \times 10^{-4}/4.1 \times 10^{-4}/4.5 \times 10^{-4}$ |
| 0.05 | 0 | 0.3 | $4.7 \times 10^{-4}/4.4 \times 10^{-4}/3.4 \times 10^{-4}$ | $2.9 \times 10^{-4}/4.2 \times 10^{-4}/3.6 \times 10^{-4}$ | $1.4 \times 10^{-3}/1.4 \times 10^{-3}/1.5 \times 10^{-3}$ | $4.7 \times 10^{-4}/4.4 \times 10^{-4}/6.0 \times 10^{-4}$ |
| 0.05 | 0 | 0.5 | $6.4 \times 10^{-4}/4.0 \times 10^{-4}/3.4 \times 10^{-4}$ | $4.9 \times 10^{-4}/5.2 \times 10^{-4}/4.8 \times 10^{-4}$ | $4.2 \times 10^{-3}/3.8 \times 10^{-3}/3.5 \times 10^{-3}$ | $4.7 \times 10^{-4}/5.0 \times 10^{-4}/4.4 \times 10^{-4}$ |
| 0.1 | 0 | 0.1 | $3.3 \times 10^{-4}/2.6 \times 10^{-4}/4.9 \times 10^{-4}$ | $5.9 \times 10^{-4}/6.2 \times 10^{-4}/4.6 \times 10^{-4}$ | $1.1 \times 10^{-3}/1.1 \times 10^{-3}/1.2 \times 10^{-3}$ | $5.3 \times 10^{-4}/5.9 \times 10^{-4}/5.3 \times 10^{-4}$ |
| 0.1 | 0 | 0.3 | $6.0 \times 10^{-4}/4.7 \times 10^{-4}/4.1 \times 10^{-4}$ | $1.6 \times 10^{-4}/1.3 \times 10^{-4}/1.3 \times 10^{-4}$ | 0.011/0.011/9.1 $\times 10^{-3}$ | $4.7 \times 10^{-4}/5.4 \times 10^{-4}/4.1 \times 10^{-4}$ |
| 0.1 | 0 | 0.5 | $6.3 \times 10^{-4}/6.7 \times 10^{-4}/6.3 \times 10^{-4}$ | $3.5 \times 10^{-4}/3.5 \times 10^{-4}/3.5 \times 10^{-4}$ | 0.038/0.040/0.032 | $5.8 \times 10^{-4}/5.9 \times 10^{-4}/4.9 \times 10^{-4}$ |
| Power |  |  |  |  |  |  |
| 0.05 | 0.1 | 0.1 | 0.21/0.21/0.19 | 0.043/0.044/0.040 | - | 0.22/0.23/0.21 |
| 0.05 | 0.1 | 0.3 | 0.19/0.19/0.17 | $1.3 \times 10^{-3}/1.3 \times 10^{-3}/1.2 \times 10^{-3}$ | - | 0.22/0.23/0.21 |
| 0.05 | 0.1 | 0.5 | 0.17/0.16/0.14 | $6.9 \times 10^{-4}/5.2 \times 10^{-4}/5.6 \times 10^{-4}$ | - | 0.22/0.23/0.21 |
| 0.1 | 0.1 | 0.1 | 0.22/0.22/0.20 | 0.048/0.048/0.043 | - | 0.22/0.23/0.21 |
| 0.1 | 0.1 | 0.3 | 0.20/0.20/0.18 | $1.1 \times 10^{-3}/1.2 \times 10^{-3}/1.1 \times 10^{-3}$ | - | 0.22/0.23/0.21 |
| 0.1 | 0.1 | 0.5 | 0.17/0.16/0.14 | $6.8 \times 10^{-4}/5.9 \times 10^{-4}/6.8 \times 10^{-4}$ | - | 0.22/0.23/0.21 |
| 0.05 | 0.2 | 0.1 | 0.59/0.60/0.58 | 0.28/0.28/0.27 | - | 0.60/0.61/0.59 |
| 0.05 | 0.2 | 0.3 | 0.57/0.57/0.55 | 0.011/0.011/0.010 | - | 0.60/0.61/0.59 |
| 0.05 | 0.2 | 0.5 | 0.54/0.53/0.52 | $4.9 \times 10^{-4}/5.9 \times 10^{-4}/6.4 \times 10^{-4}$ | - | 0.60/0.61/0.59 |
| 0.1 | 0.2 | 0.1 | 0.59/0.60/0.58 | 0.28/0.28/0.27 | - | 0.60/0.61/0.59 |
| 0.1 | 0.2 | 0.3 | 0.58/0.58/0.56 | 0.011/0.011/8.8 $\times 10^{-3}$ | - | 0.60/0.61/0.59 |
| 0.1 | 0.2 | 0.5 | 0.54/0.53/0.52 | $4.5 \times 10^{-4}/5.5 \times 10^{-4}/6.5 \times 10^{-4}$ | - | 0.60/0.61/0.59 |

\*: For the method that replaces missing summary statistics with 0, the type I error is inflated. So power was not evaluated.

Second, we found that the DISCARD method of discarding the studies with missing data produces valid type I errors, but can lead to considerable loss of power. For example, when the known variant has effect .1, the causal variant at the candidate gene has effect .2, and 30% of the contributed summary statistics in each study contain missing values, the power for DISCARD is 41%, much lower than the power for PCBS (65%). When 50% of the variant sites contain missing data, discarding studies with missing data results in even lower power (24%) compared to PCBS (64%). For gene-level association tests, discarding studies with missing summary statistics can lead to similar power loss, regardless of the rare variant association tests performed (**Table 2**).

Third, we noted that the power for conditional analysis is affected by where the missing data lies. The missing summary statistics from candidate variant sites reduce the power of single variant association tests. Yet, the PCBS statistics remained to be the most powerful.

Interestingly, gene-level association tests are affected by two types of missing data with opposite consequences: Missing values at causal variant sites reduce power but missing values at non-causal variant sites tend to reduce noise and thus improve power. The net power loss was small across all scenarios. For instance, when a causal variant in the candidate gene has effects sampled from *N*(0,0.2^2^), the conditioned variant has effect .1, and 30% of the contributed summary statistics in each study have missing values, the power for burden/SKAT/VT tests are 58%/58%/56%, which are only slightly reduced compared to the power of analyzing the complete datasets (60%/61%/60%). On the other hand, the method that discards studies with missing data had much reduced power (0.011/0.011/8.8×10^-3^).

Using PCBS (partial correlation based score statistic), the power for conditional analysis is primarily influenced by the sample size at the candidate variant site. An important observation is that when missing data only occurs at variant sites that we conditioned on, the conditional analysis using PCBS statistics of incomplete datasets attains similar power as analysis of the complete dataset (**Table 3**). The power loss is minimal even when a large number of studies contain missing summary statistics at conditioned variant sites. For example, in the scenario with known effects .1 and candidate variant effect .2, when score statistics at conditioned variant sites are missing from 50% of the studies, the power for PCBS statistics is 0.64 and the power for the analysis of the complete data is 0.66. Similar power comparisons were also observed for gene-level tests (**Table 4**).

**Table 3:** The power and type I errors for single variant conditional meta-analysis strategies in the presence of missing data. The simulation setup is the same as in Table 1, except that the missing summary statistics are only present at the conditioned variant sites.

| Conditioned Variant Effect | Candidate Variant Effect ($\tau_\beta$) | Fraction of Missing Data at Conditioned Variant Sites | Type I Error/Power | | | Analyze the Full Dataset [Gold Standard] |
| --- | --- | --- | --- | --- | --- | --- |
|  |  |  | PCBS | DISCARD | REPLACE0<sup>*</sup> |  |
| Type I Error |  |  |  |  |  |  |
| 0.05 | 0 | 0.1 | $4.3 \times 10^{-4}$ | $3.4 \times 10^{-4}$ | $5.0 \times 10^{-4}$ | $3.6 \times 10^{-4}$ |
| 0.05 | 0 | 0.3 | $3.2 \times 10^{-4}$ | $3.3 \times 10^{-4}$ | $1.9 \times 10^{-3}$ | $3.3 \times 10^{-4}$ |
| 0.05 | 0 | 0.5 | $5.1 \times 10^{-4}$ | $3.8 \times 10^{-4}$ | $6.5 \times 10^{-3}$ | $3.6 \times 10^{-4}$ |
| 0.1 | 0 | 0.1 | $4.7 \times 10^{-4}$ | $3.8 \times 10^{-4}$ | $1.3 \times 10^{-3}$ | $4.0 \times 10^{-4}$ |
| 0.1 | 0 | 0.3 | $4.0 \times 10^{-4}$ | $2.9 \times 10^{-4}$ | 0.014 | $3.9 \times 10^{-4}$ |
| 0.1 | 0 | 0.5 | $5.7 \times 10^{-4}$ | $3.6 \times 10^{-4}$ | 0.057 | $3.7 \times 10^{-4}$ |
| Power |  |  |  |  |  |  |
| 0.05 | 0.1 | 0.1 | 0.064 | 0.046 | - | 0.064 |
| 0.05 | 0.1 | 0.3 | 0.063 | 0.039 | - | 0.065 |
| 0.05 | 0.1 | 0.5 | 0.061 | 0.020 | - | 0.065 |
| 0.1 | 0.1 | 0.1 | 0.063 | 0.044 | - | 0.063 |
| 0.1 | 0.1 | 0.3 | 0.064 | 0.033 | - | 0.063 |
| 0.1 | 0.1 | 0.5 | 0.063 | 0.019 | - | 0.063 |
| 0.05 | 0.2 | 0.1 | 0.66 | 0.51 | - | 0.66 |
| 0.05 | 0.2 | 0.3 | 0.66 | 0.42 | - | 0.66 |
| 0.05 | 0.2 | 0.5 | 0.64 | 0.24 | - | 0.66 |
| 0.1 | 0.2 | 0.1 | 0.66 | 0.51 | - | 0.65 |
| 0.1 | 0.2 | 0.3 | 0.65 | 0.41 | - | 0.66 |
| 0.1 | 0.2 | 0.5 | 0.64 | 0.24 | - | 0.66 |

\*: For the method that replaces missing summary statistics with 0, the type I error is inflated. So power was not evaluated.

**Table 4:** Type I Error and Power for Gene-level Association Tests in the Presence of Missing Data. We evaluated the power for burden, SKAT and VT test in the presence of missing data. The simulation setup is the same as Table 2, except that missing summary statistics are only present in the conditioned variant site.

| Conditioned Variant Effect | Candidate Variant Effect ($\tau_\beta$) | Fraction of Missing Data at Conditioned Variant Sites | Type I Error/Power for Burden/SKAT/VT ($\alpha=0.0005$) | | | |
| --- | --- | --- | --- | --- | --- | --- |
|  |  |  | PCBS | DISCARD | REPLACE0* | Analyze the Full Dataset [Gold Standard] |
|  |  |  | Type I Error |  |  |  |
| 0.05 | 0 | 0.1 | $5.6 \times 10^{-4}/5.1 \times 10^{-4}/5.4 \times 10^{-4}$ | $5.6 \times 10^{-4}/5.3 \times 10^{-4}/4.6 \times 10^{-4}$ | $3.3 \times 10^{-4}/3.3 \times 10^{-4}/6.6 \times 10^{-4}$ | $5.3 \times 10^{-4}/5.4 \times 10^{-4}/5.6 \times 10^{-4}$ |
| 0.05 | 0 | 0.3 | $5.6 \times 10^{-4}/4.7 \times 10^{-4}/5.6 \times 10^{-4}$ | $5.2 \times 10^{-4}/5.1 \times 10^{-4}/4.7 \times 10^{-4}$ | $5.8 \times 10^{-3}/5.9 \times 10^{-3}/5.9 \times 10^{-3}$ | $4.2 \times 10^{-4}/5.4 \times 10^{-4}/5.1 \times 10^{-4}$ |
| 0.05 | 0 | 0.5 | $5.1 \times 10^{-4}/4.6 \times 10^{-4}/4.7 \times 10^{-4}$ | $5.1 \times 10^{-4}/5.2 \times 10^{-4}/4.6 \times 10^{-4}$ | $9.6 \times 10^{-3}/9.6 \times 10^{-3}/9.8 \times 10^{-3}$ | $4.9 \times 10^{-4}/5.2 \times 10^{-4}/5.7 \times 10^{-4}$ |
| 0.1 | 0 | 0.1 | $4.8 \times 10^{-4}/4.8 \times 10^{-4}/5.3 \times 10^{-4}$ | $4.8 \times 10^{-4}/4.8 \times 10^{-4}/4.5 \times 10^{-4}$ | $1.8 \times 10^{-3}/1.8 \times 10^{-3}/1.2 \times 10^{-3}$ | $7.1 \times 10^{-4}/7.1 \times 10^{-4}/5.3 \times 10^{-4}$ |
| 0.1 | 0 | 0.3 | $4.7 \times 10^{-4}/4.4 \times 10^{-4}/4.7 \times 10^{-4}$ | $4.7 \times 10^{-4}/4.7 \times 10^{-4}/5.1 \times 10^{-4}$ | 0.013/0.013/0.012 | $1.7 \times 10^{-4}/1.7 \times 10^{-4}/1.7 \times 10^{-4}$ |
| 0.1 | 0 | 0.5 | $4.9 \times 10^{-4}/4.9 \times 10^{-4}/5.6 \times 10^{-4}$ | $4.9 \times 10^{-4}/4.9 \times 10^{-4}/4.6 \times 10^{-4}$ | 0.049/0.049/0.043 | $4.9 \times 10^{-4}/4.9 \times 10^{-4}/8.2 \times 10^{-4}$ |
| Power |  |  |  |  |  |  |
| 0.05 | 0.1 | 0.1 | 0.21/0.21/0.20 | 0.19/0.20/0.18 | - | 0.22/0.22/0.21 |
| 0.05 | 0.1 | 0.3 | 0.22/0.22/0.20 | 0.15/0.15/0.14 | - | 0.22/0.23/0.21 |
| 0.05 | 0.1 | 0.5 | 0.22/0.22/0.20 | 0.090/0.091/0.084 | - | 0.23/0.23/0.21 |
| 0.05 | 0.2 | 0.1 | 0.59/0.60/0.58 | 0.57/0.57/0.56 | - | 0.60/0.61/0.59 |
| 0.05 | 0.2 | 0.3 | 0.58/0.59/0.57 | 0.49/0.50/0.48 | - | 0.59/0.60/0.59 |
| 0.05 | 0.2 | 0.5 | 0.58/0.58/0.57 | 0.39/0.40/0.38 | - | 0.59/0.60/0.58 |
| 0.1 | 0.1 | 0.1 | 0.22/0.22/0.20 | 0.20/0.21/0.19 | - | 0.23/0.23/0.21 |
| 0.1 | 0.1 | 0.3 | 0.23/0.23/0.21 | 0.15/0.16/0.14 | - | 0.24/0.24/0.22 |
| 0.1 | 0.1 | 0.5 | 0.22/0.21/0.20 | 0.090/0.089/0.081 | - | 0.23/0.23/0.21 |
| 0.1 | 0.2 | 0.1 | 0.60/0.60/0.59 | 0.58/0.59/0.57 | - | 0.60/0.61/0.60 |
| 0.1 | 0.2 | 0.3 | 0.57/0.58/0.56 | 0.49/0.49/0.48 | - | 0.58/0.59/0.58 |
| 0.1 | 0.2 | 0.5 | 0.59/0.59/0.57 | 0.41/0.41/0.39 | - | 0.60/0.60/0.59 |

We also examined if the genetic effect heterogeneity between studies would affect the performance of PCBS statistics (**S1 Table, S2 Table**). We sampled the conditional variant effect from a normal distribution, allowing the effect to vary between studies. When there was a large amount of heterogeneity in the genetic effect of the conditioned variant, the type I error remained well controlled. The power for conditional analysis appeared lower relative to the scenario where the conditioned variant had fixed effects across all studies. For example, when the candidate variant effect was 0.2, the conditioned variant effects in each cohort were sampled from *N*(0.1,0.25^2^), and the rate of missingness was 50%, the power for the conditional analysis using PCBS was 58%, slightly lower than the power when analyzing the complete dataset (67%). Yet the power for the PCBS statistic was still substantially higher than the method that discard studies with missing data (24%).

### Comparison of the Accuracy of the Genetic Effect Estimates

Finally, we evaluated the accuracy of the estimate of conditional genetic effects (**S3 Table**). We evaluated the bias and mean squared error for the three analysis strategies. The PCBS method produced unbiased estimates of conditional effects. The bias of the estimator was comparable to the method that removes studies with missing data. The PCBS method more effectively used the summary statistics across studies, and hence produced smaller mean squared error. The method that replaces missing summary statistics with zero gave highly biased estimates of conditional effect. For example, when the genetic effect of the conditioned variant was .5, and the candidate variant effect was 0.1, the bias can be as large as 0.11.

### Analysis of real data

We performed a meta-analysis of CPD phenotype in eight cohorts. The locus *CHRNA5-CHRNB4-CHRNA3* was previously identified as associated with CPD[21]. After careful quality control, 13,960 variants and 13 genes were available for analysis within the 1 million base pair window of the strongest association (15:77806023-79806023). Using the method of Li and Ji [22] that accounts for the linkage disequilibrium of tested variants, we calculated that there are the equivalent of 2452 independent tests. A significance threshold of 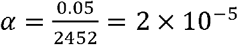 was used to identity independently associated variants.

It is important to note that even with high quality imputation panels, there is still considerable missing data in the imputed datasets. Within the locus of interest, 75.5% of the variants are missing from at least one participating studies post imputation, due to the use of different imputation panels for the UK Biobank versus the remaining studies, as well as post-imputation filtering on imputation qualities.

Using sequential forward selection with the new PCBS method, we identified three independently associated variants (rs8034191, rs3825845, rs3825930) with p-values < 2 × 10^-5^, the threshold for Bonferroni correction of testing 2,452 independently associated variants (**Table 5**). Three variants were reported to be genome-wide significant in the locus, including rs1051730, rs55958997, rs28675338. Yet, the variant rs28675338 overlaps an in-del, and thus was not included in the Haplotype Reference Consortium panel [23]. Our newly identified variants differed from previously reported top signals in the *CHRNA5-CHRNB4-CHRNA3* locus [24]. We further examined whether our top independently associated signals explained previously reported hits, by performing association analysis of previously reported variants, conditional on our top 3 independently associated variants. We noted that all of the previously reported association signals are no longer significant (p>0.05) (**S4 Table**). On the other hand, by performing conditional analysis in the opposite direction (conditional on rs1051730, rs55958997), two of our newly identified independent association (rs3825845, rs3825930) remain statistically significant, conditional on previously reported GWAS hits. We estimated the genetic variance explained by the identified independently associated variants. For the three newly identified association signals, they together explain 0.46% of the phenotypic variation. On the other hand, the known association signals (as well as their proxy in the dataset) together only explain 0.34% of the phenotypic variance. Independently associated variants detected using our new method substantially improve the phenotypic variance explained.

**Table 5:**
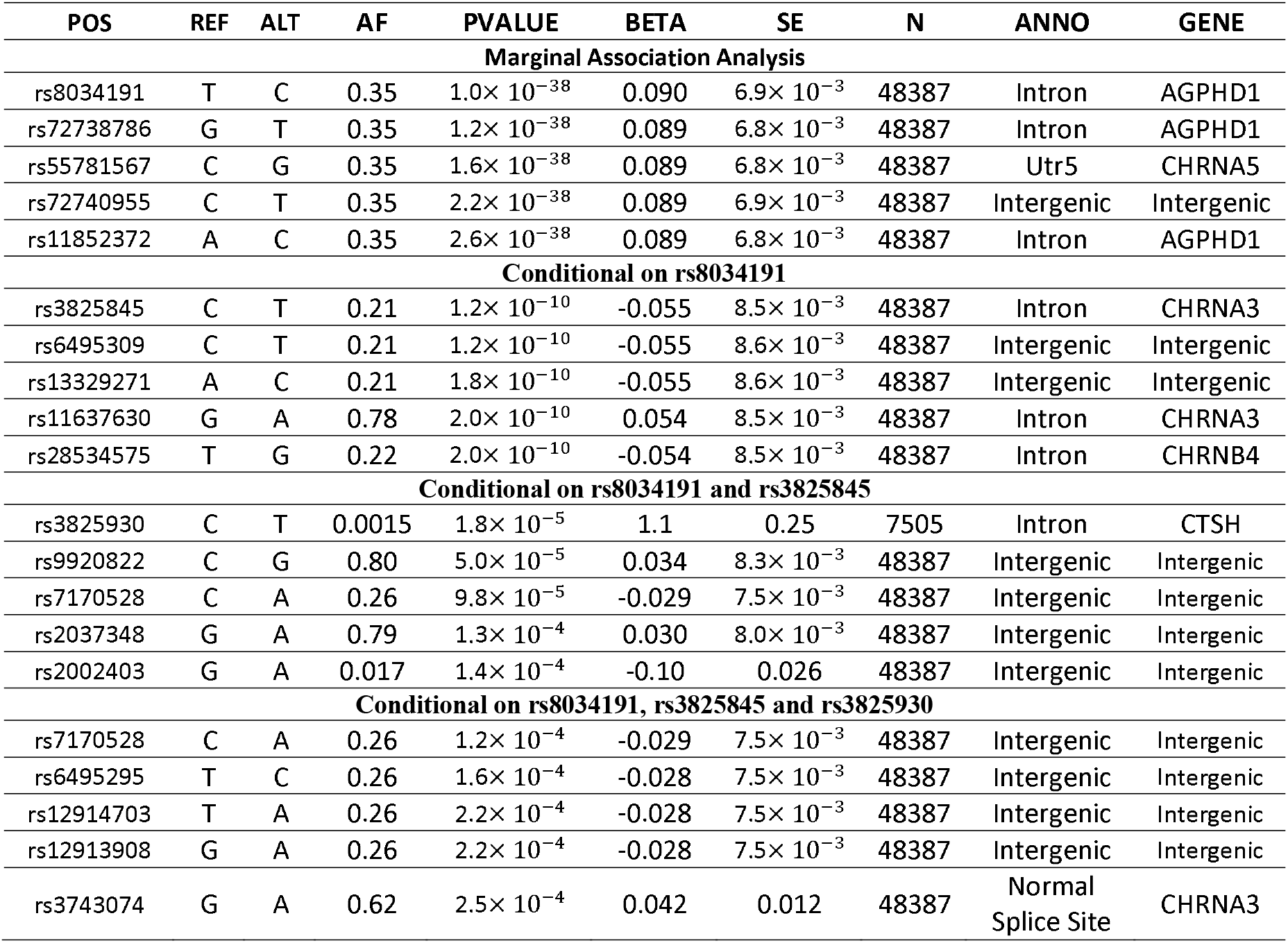
Sequential conditional analysis for the *CHRNA5-CHRNB4-CHRNA3* locus. We iteratively performed conditional analysis, conditioning on the top variants from earlier rounds. Top 5 association signals at each iteration are shown. The sequential conditional analysis stops when the top association signal is no longer significant under the Bonferroni correction threshold *α* = 2 × 10^−5^.

As a comparison, we also performed sequential forward selection using the two alternative approaches. Using the DISCARD method, no additional association signals are identified beyond the top association signal. Using REPLACE0, only two independently associated variants were identified, i.e. rs8034191, rs3825845. Both REPLACE0 and DISCARD failed to identify rs3825930. Concordant with our simulation study, the result of PCBS statistics differ from REPLACE0 and DISCARD, where a large number of missing values are present in the contributed summary association statistics (**S5 Table**).

Finally, we asked if rare variants within the *CHRNA5-CHRNB4-CHRNA3* locus are independently associated with the CPD phenotype (S6 Table). Thirteen genes were analyzed using simple burden, SKAT and VT tests under a MAF threshold of 0.05. None of the resulting p-values were less than 0.05/13.

## DISCUSSION

We proposed a simple yet effective meta-analysis method to estimate joint and conditional effects of rare variants in the presence of missing summary statistics from contributing studies. The method leads to the optimal use of shared summary association statistics. It has well controlled type I error and much higher power than alternative approaches even when a large number of contributing studies contain missing summary statistics.

A tempting alternative to using partial correlation based score statistics is to impute missing summary association statistics before meta-analysis. Recently, Gaussian imputation methods[25-27] were developed to directly impute summary association statistics without resorting to individual-level data. However, Gaussian imputation shares similar issues with hidden Markov model based methods (e.g. it cannot impute well for studies that use targeted genotyping or sequencing assays). Imputing low frequency variants is also challenging. As such, it is often recommended to discard imputed summary statistics for low frequency variants. Our proposed method (PCBS) can nicely complement imputation-based methods when accurate imputation is infeasible. It is also important to note that our method is not a replacement of imputation methods. Imputation methods, if feasible, increase effective sample sizes for imputed variants, and increase power. Our method, on the other hand, does not increase the effective sample size for tested variants. In practice, imputation method should first be applied in each participating cohort. Our method should be applied at the meta-analysis stage for valid and powerful conditional meta-analysis, especially when contributed summary statistics from participating cohorts contain missing values.

Missing data will continue to be a persistent issue in the next generation of large-scale genetic studies. Major biobanks have started to develop their own genotyping arrays and imputation reference panels to incorporate customized content. Combining these newly genotyped studies with existing datasets will result in missing summary statistics. Our method will continue to be useful when analyzing these newly generated datasets.

Another major application of the proposed method is in the meta-analysis of sequence data. Given the use of targeted sequencing assays and variability in batch processing and quality control across studies, it would be difficult to impute missing genotype data or missing summary statistics. One of the challenges in sequence-based meta-analysis is to properly represent monomorphic sites, as the polymorphic variant sites are not known a priori. Neither un-called variant sites (e.g. due to insufficient coverage or failed quality control) nor monomorphic sites contribute to the single variant meta-analysis statistic. Yet they should be treated differently in joint and conditional meta-analysis. Summary statistics from monomorphic variants should be replaced by zero. On the other hand, summary statistics from un-called variants should be treated as missing data, and the conditional association analysis can be performed using our partial correlation based score statistics.

While not the focus of this article, the proposed method is also helpful for downstream analyses that make use of the joint effects of multiple variants, e.g. estimating the phenotypic variance explained by independently associated variants. The validity of these analyses critically rely on the proper estimates of joint effects, which are usually obtained from single variant association statistics and the LD information from a reference panel. When summary statistics from contributing studies contain missing data, the correlations between resulting marginal meta-analysis association statistics may not be properly approximated by the R^2^ estimated from a reference panel. In this case, PCBS can be used to obtain valid joint effect estimates, which can potentially lead to better calibrated phenotypic variance explained.

Our paper focused on exact conditional analysis, which relies on the exact covariance matrices of score statistics shared across studies. We did not consider approximate conditional analysis that makes use of LD matrices from reference panels to approximate the covariance between score statistics[28]. It was shown that approximate conditional analysis can be less accurate than the exact methods for rare variant association studies [29]. In the presence of missing summary statistics from contributing studies, the approximate conditional analysis method may often incorrectly estimate covariance matrices between score statistics. For example, consider a simple example of meta-analysis of two studies of equal size N. For a genetic variant that is only measured in study 1 and a genetic variant that is only measured in study 2, the resulting meta-analysis score statistics from the two sites are uncorrelated. The approximate conditional analysis may incorrectly estimate the correlation by the LD (or a scaled version of LD) between the two variant sites, which can result in invalid association analysis results. When summary statistics from contributing studies are available, we can approximate the score statistics and covariance matrix using the genetic effect estimates, their standard deviation as well as the LD information from a reference panel[29]. Our proposed methods can thus be adapted in approximate conditional analysis to obtain valid results in the presence of missing values from contributed summary statistics.

Taken together, our partial correlation based score statistic is a simple yet effective method for estimating joint and conditional effects from meta-analysis. With its efficient implementations in RVTESTS and RAREMETAL, these methods will have broad application in current array-based meta-analysis, as well as the upcoming haplotype reference consortium imputation-based meta-analysis and sequence-based meta-analysis. Correct inference on the joint and conditional effects using these methods will pave the way for a more accurate characterization and a more complete understanding of the genetic architecture for complex traits.

## Acknowledgements

This research has been conducted using the UK Biobank Resource. DJL, DM and YJ were supported by R01HG008983. MB was supported by R01HG000376, W.G.I has been supported by DA05147 and DA036216 from the National Institute of Health. SML was supported by K01HL125858. CADD study has been supported by DA011015 from the National Institute of Health. The COPDGene Study (NCT00608764) is supported by National Heart, Lung and Blood Institute NHLBI R01 HL084323 and HL089897 and is also supported by the COPD Foundation through contributions made to an Industry Advisory Board comprised of AstraZeneca, Boehringer Ingelheim, Novartis, Pfizer, GlaxoSmithKline, Siemens, and Sunovion. The funding sources played no role in the design of the study.

## Supporting Information Legends

**S1 Text.**

**S1 Table: Power and Type I Errors of Meta-analysis of Single Variant Tests in the Presence of Missing Data and Genetic Effect Heterogeneity**. We evaluated the impact of large genetic effect heterogeneity on the power and type I errors for the PCBS statistics. The effects of the conditioned variants in each cohort are sampled from the distribution *N*(*μ*_*β*_2__, 0.25^2^). All other simulation settings are the same as in Table 1.

**S2 Table: Power and Type I Errors of Meta-analysis of Gene-level Tests in the Presence of Missing Data and Genetic Effect Heterogeneity**. We evaluated the impact of large genetic effect heterogeneity on the power and type I errors for the PCBS statistics. The genetic effects for the conditioned variants in each cohort are sampled from the distribution *N*(*μ*_*β*_2__, 0.25^2^). All other simulation settings are the same as in Table 2.

**S3 Table: Accuracy of Estimates of Conditional Effects**. We compared the accuracy of the estimated genetic effects of candidate variants conditioning on 3 randomly chosen variants with effect 0.1. The absolute bias and the mean squared error for the candidate variant conditional effect estimate are displayed for different combinations of the candidate genetic variant effects and the fraction of missing data at conditioned variant site.

**S4 Table: Two Way Conditional analysis of Independently Associated Variants and Previously Reported GWAS Hits**.

**S5 Table: Results of Sequential Forward Selection Using the Method that Replaces Missing Data with 0 (Panel A), and the Method that Discards Studies with Missing Data (Panel B)**

**S6 Table: Gene-level Conditional Analysis Results**. We analyzed gene-level association test conditional on the three independently associated variants (i.e. rs8034191, rs3825845 and rs3825930), which were identified using sequential forward selection. Three gene level association tests were performed, including simple burden tests, SKAT and VT. No significant gene-level associations were identified (p<0.05/13)

